# Effects of Variation in Urine Sample Storage Conditions on 16S Urogenital Microbiome Analyses

**DOI:** 10.1101/2022.07.05.498913

**Authors:** Tanya Kumar, MacKenzie Bryant, Kalen Cantrell, Se Jin Song, Daniel McDonald, Helena M. Tubb, Sawyer Farmer, Emily S. Lukacz, Linda Brubaker, Rob Knight

**Affiliations:** Medical Scientist Training Program, University of California San Diego, La Jolla, CA, USA; Department of Pediatrics, University of California San Diego, La Jolla, CA; Department of Computer Science and Engineering, University of California San Diego, La Jolla, CA, USA; Center for Microbiome Innovation, Jacobs School of Engineering, University of California San Diego, La Jolla, CA, USA; Department of Obstetrics, Gynecology and Reproductive Sciences, University of California San Diego, La Jolla, CA, USA; Department of Bioengineering, University of California San Diego, La Jolla, CA, USA

## Abstract

Replicability is a well-established challenge in microbiome research with a variety of contributing factors at all stages, from sample collection to code execution. Here, we focus on voided urine sample storage conditions for urogenital microbiome analysis. Using urine samples collected from 10 healthy adult women, we investigated the microbiome preservation efficacy of AssayAssure® Genelock (as opposed to no preservative) under different temperature conditions. We varied temperature over 48 hours in order to examine the impact of conditions samples may experience with home voided urine collection and shipping to a central biorepository. The following common lab and shipping conditions were investigated: -20C, ambient temperature, 4C, a freeze-thaw cycle, and a heat cycle. At 48 hours, all samples were stored at -80C until processing. After generating 16S rRNA gene amplicon sequencing data using the highly sensitive KatharoSeq protocol, we observed individual variation in both alpha and beta diversity metrics below interhuman differences, corroborating reports of individual microbiome variability in other specimen types. While there was no significant difference in beta diversity when comparing AssayAssure® Genelock vs. no preservative, we did observe a higher concordance with AssayAssure samples shipped at colder temperatures (−20C and 4C) when compared to the samples shipped at -20C without preservative. Our results indicate that AssayAssure does not introduce a significant amount of microbial bias when used on a range of temperatures but is most effective at colder temperatures.

**Importance:** The urogenital microbiome is an understudied yet important human microbiome niche. Research has been stimulated by the relatively recent discovery that urine is not sterile: urinary tract microbes have been linked to health problems including urinary infections, incontinence, and cancer. The quality of life and economic impact of UTIs and urgency incontinence alone are enormous, with $3.5 billion and $82.6 billion respectively spent in the U.S. annually. Given the low biomass of urine, novelty of the field, and well-established replicability bias in microbiome studies, it is critical to study storage conditions on urine samples to minimize microbial biases. Efficient and reliable preservation methods permit home self-sample collection and shipping, increasing the accessibility of larger-scale studies. Here, we examined both buffer and temperature variation effects on 16S rRNA gene amplicon sequencing results from urogenital samples, providing data on the consequences of common storage methods on urogenital microbiome results.

## Observation

The urogenital microbiome is a relatively understudied component of the human microbiome. Additional research is warranted as it has biologic plausibility in urinary tract health and disease. It has been tied to some of the most common urinary conditions: lower urinary tract symptoms (LUTS), which include incontinence and urinary tract infections (UTIs). UTIs impact over 150 million people per year globally with many experiencing recurrent UTIs resistant to treatment, increasing the risk of serious complications (1). Thus, an in-depth understanding of the urogenital microbiome through large-scale studies is necessary to better understand urinary conditions and consider novel therapeutics.

Key advances in the field have been made recently, stimulated by the discovery that the bladder is not sterile (2, 3). Given the rapidly increasing interest in the urogenital microbiome and established replicability issues with respect to storage conditions in fecal microbiome studies, it is critical to investigate the effects of various storage conditions on urine samples (2, 4, 5). By establishing best practices for urine storage with respect to microbiome studies, we aim to lay the groundwork for population-level studies.

In this study, we examined the impact of varying temperatures on the diversity characteristics of voided urine samples in the presence of a clinically useful nucleic acid preservative, as described in **Figure 1A**. Voided urine samples, as opposed to catheterized collection, represent the urogenital microbiome (6). All samples were collected at room temperature and, informed by the work of Song et al. (7) and Marotz et al (8), immediately stored in one of five common shipping and lab conditions for 48 hours: -20C (samples were immediately frozen at -20C after transportation on dry ice), 4C (samples were transported and kept at 4C), ambient temperature (samples were transported and kept at ambient temperature), heat cycle (samples were alternated between 4C and 40C at 12 hour intervals), and freeze-thaw cycle (samples were alternated between -20C and ambient temperature at 12 hour intervals). After 48 hours, all samples were kept at -80C until processing.

**Figure 1.**
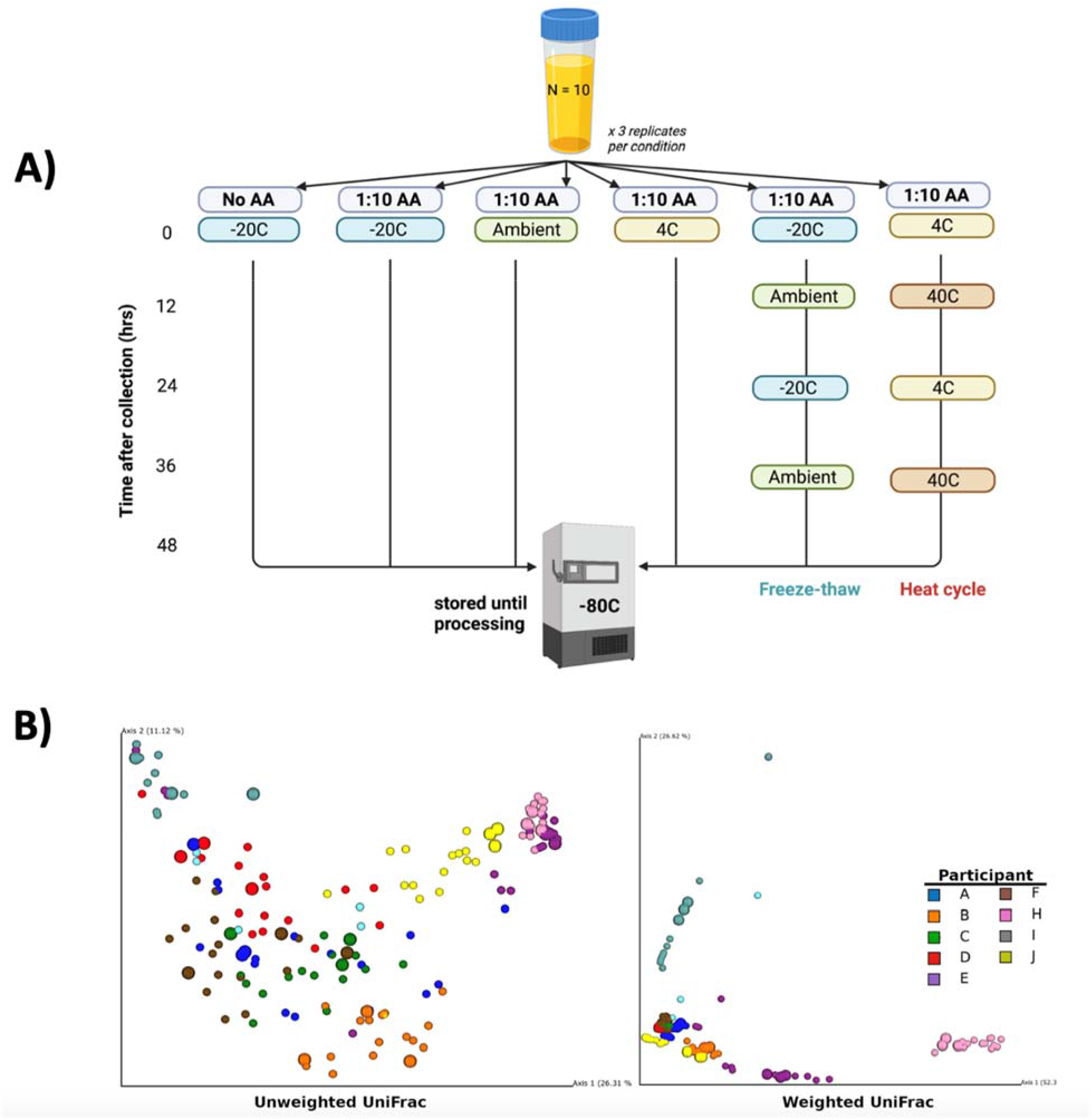
Experimental Design and Clustering by Individual. (A) Ten healthy adult women gave a single 10 mL urine sample, which was processed in triplicate and subject to varying temperature conditions and the presence or absence of AssayAssure® Genelock (AA). This figure was created with BioRender.com. (B) Principal-coordinate analysis plots of weighted and unweighted UniFrac distances.

Using the temperature conditions above, we evaluated the efficacy of AssayAssure® Genelock, a nucleic acid preservative formulated by Sierra Molecular for clinical urogenital sample collection (9–12). Jung et al. (13) found temperature-dependent biases when examining AssayAssure® Genelock on urine samples at -20C, 4C, and room temperature, prompting us to further investigate with an extended list of temperature conditions.

Following IRB approval (UCSD IRB protocol #801735) and with verbal consent, 10 healthy adult women donated a single 10mL urine sample using the Peezy Midstream (Forte Medical) collection device in a clinical setting. Urine aliquots of 500uL were immediately transferred from the Peezy tube into 15mL screw cap tubes (Sarstedt, Inc.) containing 1:10 AssayAssure® Genelock (Sierra Molecular) or no AssayAssure®. All samples included 3 technical replicates. Immediately after aliquoting the samples, the samples were stored in their respective temperature condition and transported to the lab for storage and processing.

Samples were processed using Earth Microbiome Project standard protocols (https://earthmicrobiome.org/protocols-and-standards/), as updated in Shaffer, et al. (14). Urine samples and serially diluted positive positive controls (*Paracoccus denitrificans* and *Bacillus subtilis*) were plated into 96-well MagMAX Microbiome Bead Plates (ThermoFisher Scientific) and extracted using the MagMAX Microbiome Ultra NA Extraction Kit (ThermoFisher Scientific).

Positive controls were used to account for sequencing difficulty often encountered with low biomass samples, such as urine, due to potential trace contamination from DNA extraction or PCR kit reagents. The highly sensitive KatharoSeq protocol was used to overcome this issue by using serially diluted bacterial mock community controls to utilize known read counts as a sample exclusion threshold, or limit of detection (15).

After extraction, the 16S rRNA V4 region was amplified with unique Golay barcodes on the 515f forward primer to allow post-sequencing demultiplexing. Low-cost miniaturized amplicon PCR reactions were used, as outlined in Minich, et al (16). Post-PCR 16S rRNA libraries were equal volume pooled and sequenced on a MiSeq (Illumina).

Forward read sequences generated from the MiSeq were trimmed to 150 nucleotides, quality filtered, and demultiplexed using Qiita (Qiita study 14383, EBI accession ERP138439) (17). We utilized the 50% KatharoSeq threshold to exclude and rarefy samples to 986 reads, resulting in a final analysis pool of 9 participants.

We first examined the beta diversity metrics of the samples (**Figure 1B**). In both weighted and unweighted UniFrac, permutational multivariate analysis of variance (PERMANOVA) revealed that beta diversity was driven primarily by participant (PERMANOVA, unweighted p=0.001, f=17.85; weighted p=0.001, f=54.5), rather than preservative method (PERMANOVA, unweighted p=0.96, f=0.46; weighted p=0.43, f=0.87) or temperature treatment (PERMANOVA, unweighted p=0.76, f=0.84; weighted p=0.25, f=1.22). This result is in line with previous reports on other microbiome sites (8, 13), which indicate an individual’s microbiome composition accounts for a large amount of the variation in beta diversity. We also observed that weighted UniFrac showed a higher degree of clustering than unweighted.

Next, we examined the effect of temperature by comparing the UniFrac distances between the different temperature treatment groups and samples stored immediately at -20C without AssayAssure® Genelock, our baseline group. We observed that this treatment tends to be below the mean distance between participants (interhuman), suggesting that the host is the primary driver of beta diversity. Additionally, in the case of unweighted UniFrac, the distance is close to the mean distance between replicates, suggesting the temperature plays a minor role in the microbial composition (**Figure 2A)**. The mean differences in both alpha diversity measures (Shannon and Faith PD) were close to 0 (**Figure 2B**). Thus, the host is a major contributor for both microbial community composition (**Figure 2A)** and diversity (**Figure 2B**). **Figure 2B** also depicts individual diversity variations, with some participants (such as participant B) having higher deviations from the baseline, while others (such as participant F) having an overall lower reading on richness, evenness, and phylogenetic-based diversity. We compared the relative abundance of each genus in each participant’s sample processed with a temperature treatment to their corresponding baseline (**Figure 2C)**. Similar to Song et al. (7), cold temperatures (such as -20C and 4C) had very little change in relative abundance (correlation coefficient, *r* ≥ 0.97) while freeze-thaw and heat temperatures (correlation coefficient, r ≥ 0.9) resulted in a small shift in abundance. However, ambient temperatures resulted in a large shift in relative abundance (correlation coefficient, r ≥ 0.82).

**Figure 2.**
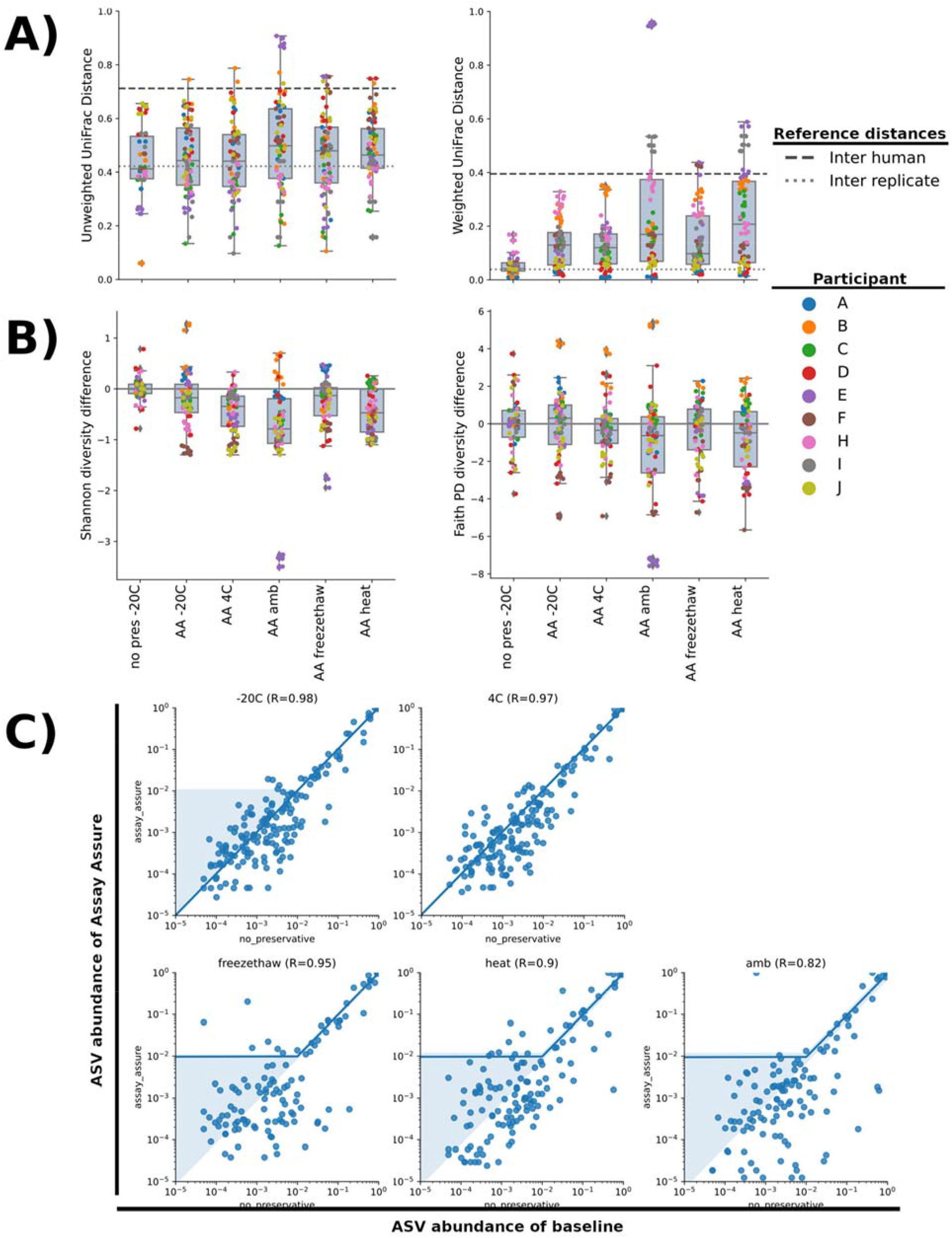
Quantification of Results. Effect of temperature treatment and presence of AssayAssure® Genelock on urine microbiome composition. AssayAssure® Genelock is present in all samples other than baseline, which was stored at -20C with no preservative (no pres). (A) Unweighted and weighted UniFrac distances between different temperature treatments and baseline. (B) Shan non and faith PD alpha diversity differences between different temperature treatments and baseline. (C) Pearson correlation where each point represents a genus with relative abundance in the baseline samples on the x-axis and the relative abundance in the different temperature treatment groups on the y-axis. Both axes are presented in log base 10 scale.

Overall, our results support the growing observation that the urogenital microbiome is a driver of inter-individual microbial diversity and should be taken into account in personalized healthcare. Our data indicate that AssayAssure® Genelock does not introduce more microbial bias in urine samples than differences attributable to individual variation while sampling across a variety of commonly found temperatures in lab and shipping settings. Thus, it is acceptable as a urine preservative for use in microbiome studies. The most minimal disruption of microbial communities occurs in cold temperature storage. These data are important to note when designing population-based studies, as current methods for urine shipping and preservation with AssayAssure® Genelock are sufficient for urogenital microbiome studies, thus allowing for participant self-collection. Future large-scale studies are warranted to understand individual urogenital microbiomes and their relationship to urinary tract health, and self-collection can increase participant accessibility.

## Acknowledgments

We thank Karenina Sanders for help in formatting citations and paper submission, and Gail Ackermann for education and help in initial Qiita data processing.

## Funding information

This work was supported by the following grants: NIH T32 GM719876, NIH/NIDDK 1U01DK106827.

## Competing Interests

None

## Data availability

A STORMS (Strengthening The Organizing and Reporting of Microbiome Studies) checklist (18) is available at [10.5281/zenodo.6788075].

